# Wide-Scale Comparative Analysis of Longevity Genes and Genetic Interventions

**DOI:** 10.1101/104182

**Authors:** Hagai Yanai, Arie Budovsky, Thomer Barzilay, Robi Tacutu, Vadim E. Fraifeld

## Abstract

Hundreds of genes have been identified as being involved in the control of lifespan in the four common model organisms (yeast, worm, fruit fly and mouse). A major challenge is to determine if longevity-associated genes (LAGs) are model-specific or may play a universal role as longevity regulators across diverse taxa. A wide-scale comparative analysis of the 1,805 known LAGs across 205 species revealed that (i) LAG orthologs are substantially over-represented, from bacteria to mammals, especially noted for essential LAGs; (ii) the effects on lifespan, when manipulating orthologous LAGs in different model organisms, were mostly concordant, despite of a high evolutionary distance between them; (iii) the most conserved LAGs were enriched in translational processes, energy metabolism, development, and DNA repair. The least conserved LAGs were enriched in autophagy (Fungi), G-proteins (Nematodes), and neuroactive ligand-receptor interactions (Chordata). The results also suggest that antagonistic pleiotropy is a conserved principle of aging.

## Background

The role of genetic factors in determination of longevity and aging patterns is an intensively studied issue (Vijg and Suh 2005; Kenyon 2010). Hundreds of genes have thus far been identified as being involved in the control of lifespan in model organisms (Tacutu et al. 2013). These genes (further denoted as longevity-associated genes, LAGs) could be defined as genes whose modulation of function or expression (such as gene knockout, overexpression, partial or full loss-of-function mutations, RNA interference or genetic polymorphisms) results in noticeable changes in longevity or the aging phenotype (Budovsky et al. 2007; Tacutu et al. 2013). We have previously investigated the characteristic features of LAGs and found that (i) they display a marked diversity in their basic function and primary cellular location of the encoded proteins (Budovsky et al. 2007); (ii) LAG-encoded proteins display a high connectivity and interconnectivity. As a result, they form a scale-free protein-protein interaction network (“longevity network”), indicating that LAGs could act in a cooperative manner (Budovsky et al. 2007; Wolfson et al. 2009; Tacutu et al. 2010, 2012). (iii) Many LAGs, particularly those that are hubs in the “longevity network”, are involved in age-related diseases (including atherosclerosis, type 2 diabetes, cancer, and Alzheimer’s disease), and in aging-associated conditions (such as oxidative stress, chronic inflammation, and cellular senescence) (Budovsky et al. 2007, Budovsky et al. 2009; Tacutu et al. 2010, 2011). (iv) The majority of LAGs established by that time in yeast, worms, flies, and mice, have human orthologs, indicating their conservation “from yeast to humans” (Budovsky et al. 2007, Budovsky et al. 2009). This assumption was also supported by studies on specific LAGs or pathways such as Foxo (Martins et al. 2016), insulin/IGF1/mTOR signaling (Tatar et al. 2003; Warner 2005; Piper et al. 2008; Ziv and Hu 2011; Gems and Partridge 2013; Zhang and Liu 2014; Pitt and Kaeberlein 2015), Gadd45 (Moskalev et al. 2012), and cell–cell and cell–extracellular matrix interaction proteins (Wolfson et al. 2009). Again, the above studies were limited only to the four model organisms and humans.

Now, the existing databases on orthologs, allow for an essential extension of the analysis of LAG orthology, far beyond the traditional model organisms and humans. In particular, the data deposited in the InParanoid database – Eukaryotic Ortholog Groups (http://inparanoid.sbc.su.se/, (Sonnhammer et al. 2015)) includes orthologs for the complete proteomes of 273 species. Here, we report the results of an unprecedentedly wide-scale analysis of 1,805 LAGs established in model organisms (available at Human Ageing Genomic Resources (HAGR) – GenAge database. http://genomics.senescence.info/genes/longevity.html, (Tacutu et al. 2013)), with regard to their putative relevance to public and private mechanisms of aging.

## Results

### Orthology of longevity-associated genes

Our first question was how LAGs orthologs are distributed across diverse taxonomic groups. For that purpose, we extracted the LAG orthologs for all the species in the InParanoid database, using a software developed in our lab (see Methods). For each gene of interest, the evolutionary conservation was evaluated as the presence or absence of orthologs across 205 proteomes (all species available excluding parasites) for a high InPanaranoid score of 1.0. Parasites were excluded from the analysis because they usually keep the minimal set of genes required for survival in the hosts and thus their inclusion could bias the results into overstating the conservation of these genes and diminish the conservation of others.

As seen in Fig. 1, for the vast majority of InParanoid species, the fraction of conserved genes was significantly higher for LAGs than for the entire proteome of the same model organism. The few exceptions were fringe cases where the baseline orthology is either very high (phylogenetically very close species, for example, *C. elegans* and *C. briggsae*), or very low (phylogenetically very distant species, for example, *M. musculus* and *K. cryptofilum*) (Suppl. Table 1).

**Fig. 1.**
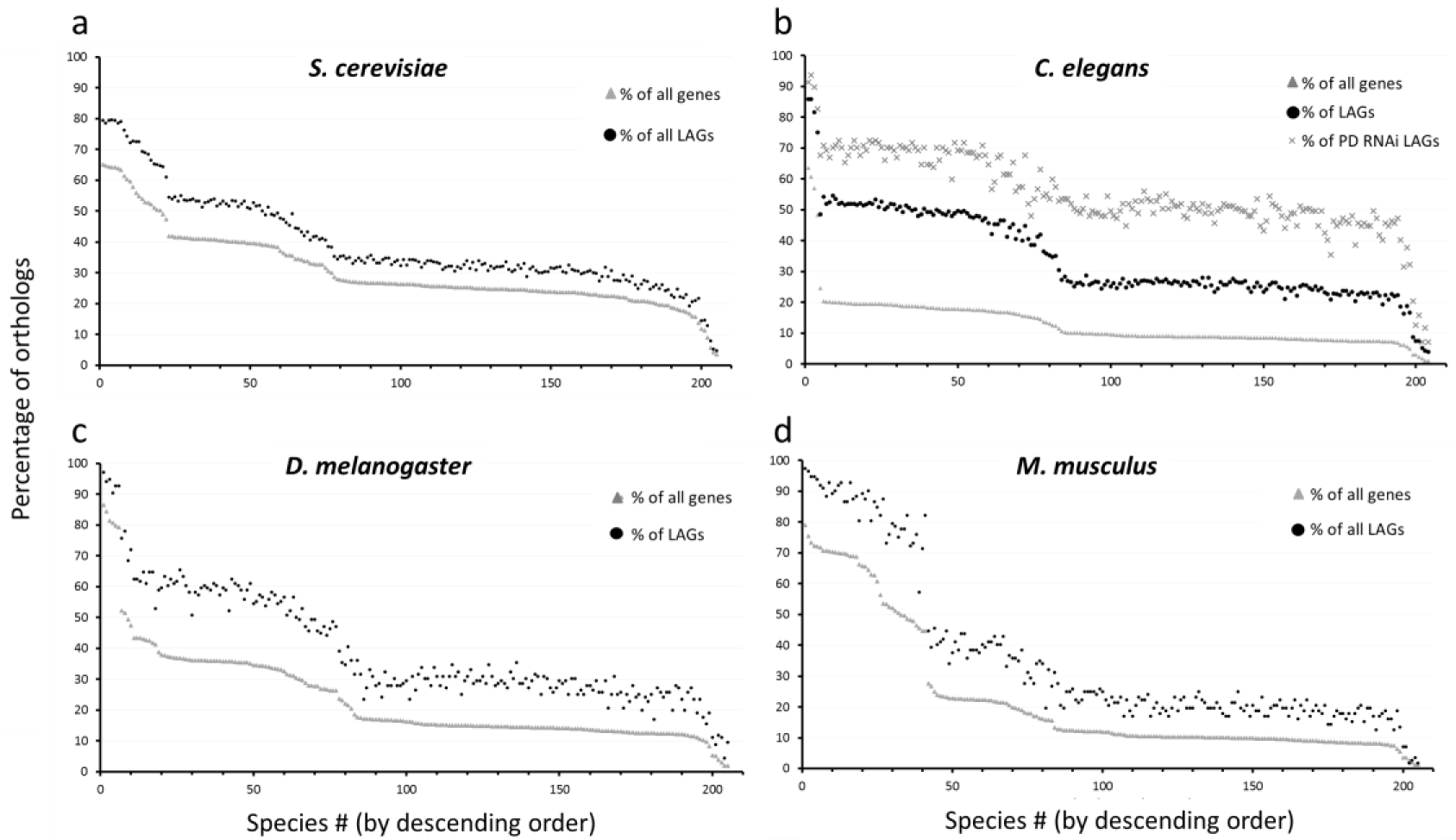
Percentage of orthologs of longevity-associated genes (LAGs) from the four model organisms across 205 species. Each graph represents one of the four model organisms and the LAGs discovered for that species. Each dot represents the percentage of orthologs between the model species and a single other species (total of 205 species from all Kingdoms, for a full list of species see Suppl. Table 1). The entire proteome of the model species (extracted from the InParanoid database) was used as control. The species (X-axis) are ordered in descending order of orthology percentage for the entire proteome. Presented are the ortholog percentage of LAGs (black circle), entire proteome (grey triangle), and *C. elegans* essential LAGs discovered by post developmental RNAi (grey x). (**a)** *S. cerevisiae*, n = 6,590 for control and 824 for LAGs; (**b)** *C. elegans*, n = 20,325 for control and 733 for LAGs; (**c)** *D. melanogaster*, n = 13,250 for control and 136 for LAGs; (**d)** *M. musculus*, n = 21,895 for control and 112 for LAGs. The vast majority of pair-wise differences between LAGs and the entire proteome are significant (p < 0.05), with a few exceptions of fringe cases as described in the text.

Remarkably, despite the high diversity of the species under analysis, the ratio between the LAG orthologs and the orthologs of the entire proteome was relatively constant along the evolutionary axis (Fig. 2). This could indicate that the high conservation of LAGs is relatively independent of evolutionary distance.

**Fig. 2.**
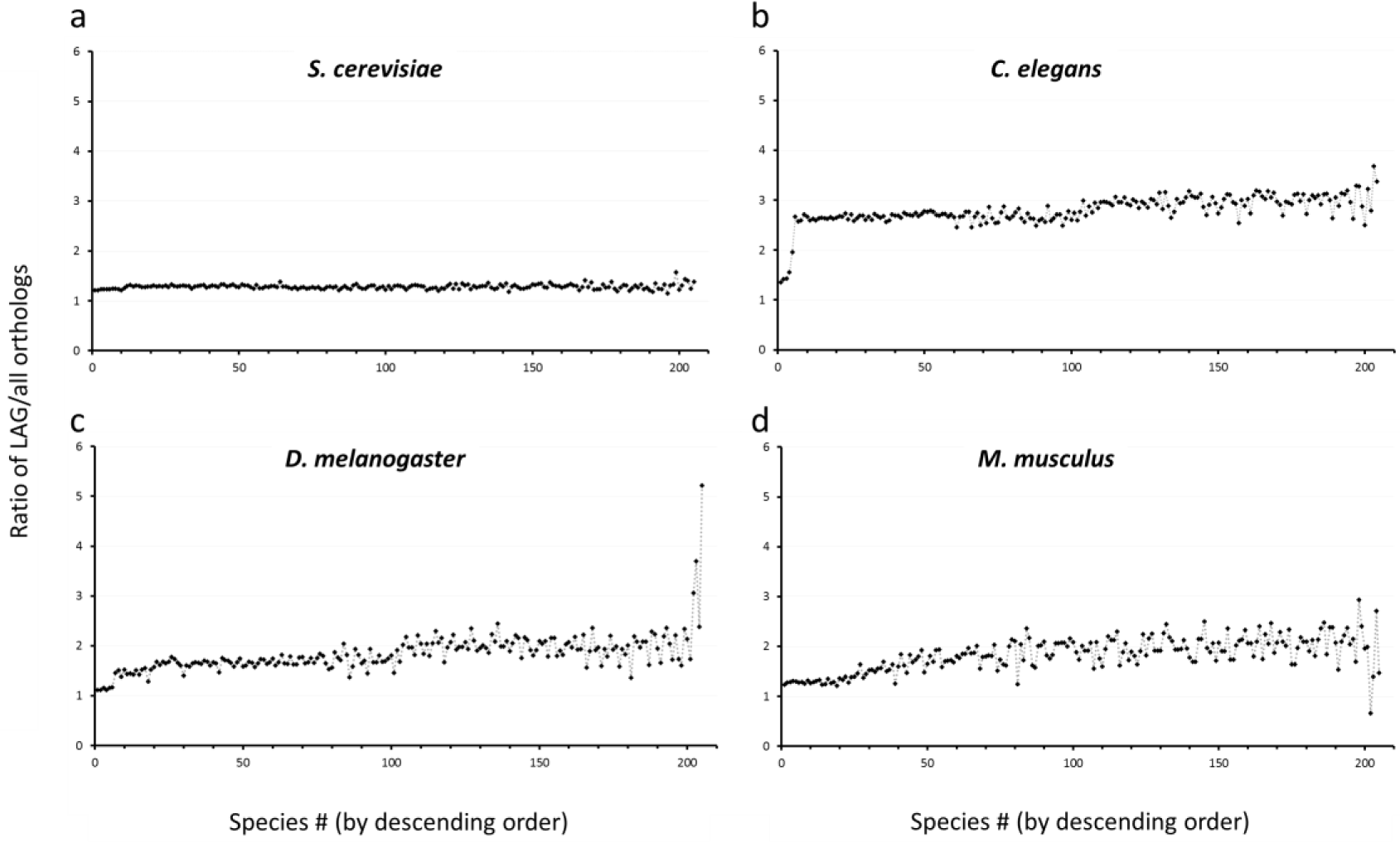
Ratio of LAGs orthologs to the entire proteome. Each graph represents the LAGs discovered in the indicated model species. Each dot represents the ratio between the orthology for LAGs to the entire proteome, for a single other species (total of 205 species from all Kingdoms, for a full list of species see Suppl. Table 1). The species (X-axis) are ordered in descending order of orthology percentage for the entire proteome. (**a)** *S. cerevisiae*, n = 6,590 for control and 824 for LAGs; (**b)** *C. elegans*, n = 20,325 for control and 733 for LAGs; (**c)** *D. melanogaster*, n = 13,250 for control and 136 for LAGs; (**d)** *M. musculus*, n = 21,895 for control and 112 for LAGs.

It should be taken into account that genes that *a priori* have orthologs in humans and are involved in basic biological processes or major diseases are more often tested for their potential effect on lifespan. Despite this obvious bias, an important observation is that among the model organisms examined, the highest conservation ratio was observed for *C. elegans* (p < E-8 for all comparisons; Fig. 2a), where the majority of LAGs were identified by means of an unbiased genome-wide RNA interference (RNAi) screens (Lee et al. 2003; Hamilton et al. 2005; Hansen et al. 2005; Yanos et al. 2012).

Using post developmental interventions such as RNAi has allowed for discovering worm longevity regulators that could not be discovered otherwise, because their pre-developmental inactivation causes a lethal phenotype (Tacutu et al. 2012). According to WormBase (http://www.wormbase.org/ (Howe et al. 2016)), 127 of the 733 known *C. elegans* LAGs are essential for development and growth, meaning an enrichment by approximately 5-fold compared to the entire genome (Tacutu et al. 2012). This is even more pronounced among LAGs that extend worm lifespan by more than 20% when inactivated: they are enriched 15-fold in essential genes. Since essential genes are generally more evolutionary conserved than non-essential ones (Tacutu et al. 2011), we compared the evolutionary conservation of the essential worm LAGs to all LAGs and found that these 127 essential LAGs are dramatically more conserved (Fig. 1a). As these genes are essential for the early stages of life, but obviously have detrimental effects later in life they, by definition, fit well into Williams’s idea of antagonistic pleiotropy (Williams 1957). If so, the results suggest that antagonistic pleiotropy is a conserved principle of aging.

One of the strong features of InParanoid is that it provides the best balance between sensitivity and specificity (Chen et al. 2007). Yet, the proteomes found in the InParanoid database contain many poorly annotated proteins and predicted transcripts that were not experimentally verified (Sonnhammer et al. 2015). These proteins have relatively few orthologs in other species and therefore could influence the results. In contrast, the interactomes from the BioGrid database (http://www.thebiogrid.org) almost exclusively include experimentally verified proteins (Chatr-Aryamontri et al. 2015). Therefore, the BioGrid data could serve as an additional, high quality control for a more rigorous testing of the evolutionary conservation of LAGs. For this purpose, we used the interactomes of *S. cerevisae*, *C. elegans*, and *D. melanogaster*. As seen in Suppl. Fig. 1, the same trend of over-conservation of LAGs was also observed in comparison to the BioGrid control. Mouse was not included in the analysis because its BioGrid gene list still contains a relatively small portion of the entire genome and thus could not provide a reliable control.

Altogether, the results clearly show a high evolutionary conservation of LAGs across distant species. With regard to this, a question arises as to whether this observation is attributed to an enrichment of specific categories that are known to be strongly preserved in the course of evolution. From the available data on gene and protein annotations for the four model species we noted that LAGs are enriched in genes that belong to categories known to be extraordinarily conserved in evolution, such as the ribosomal or mitochondrial genes (Suppl. Tables 2 and 3). However, exclusion of LAGs belonging to these categories from the analysis had almost no impact (Suppl. Fig. 2). Therefore, we conclude that the high evolutionary conservation of LAGs is not solely attributed to an enrichment of proteins from exceptionally conserved categories, but rather reflects a general trend.

### “Public” and “private” LAG categories

The distinction between public and private mechanisms of aging and longevity is a fundamental question in comparative studies of biogerontology (Gems and Partridge 2013). We attempted to shed some light on this subject based on the evolutionary conservation of LAGs in different taxa. Yet, it is important to note that if a given LAG is highly evolutionary conserved, it does not automatically translate to its role in a public mechanism of aging. In fact, in order to draw conclusions on public or private mechanisms from the presence or lack of orthologs, one must (i) have a context on the mode of operation of a given protein as its function could differ between species; or (ii) compare groups of proteins belonging to a certain pathway or category, so that generalized assumptions may be made. The data used in this study only allows for the second approach. Thus, we comprised lists of proteins under different conservation criteria, e.g. proteins that have orthologs across at least 12 phyla or have orthologs in a limited number of taxa only (for more details, see Suppl. Table 4). As shown in Fig. 3, LAGs are generally conserved over more phyla than the entire genome, again indicating their wider evolutionary conservation. Nevertheless, while most LAGs are broadly conserved, a considerable portion of them (around 10-20%) are specific to a relatively small number of phyla (Fig. 3).

**Fig. 3.**
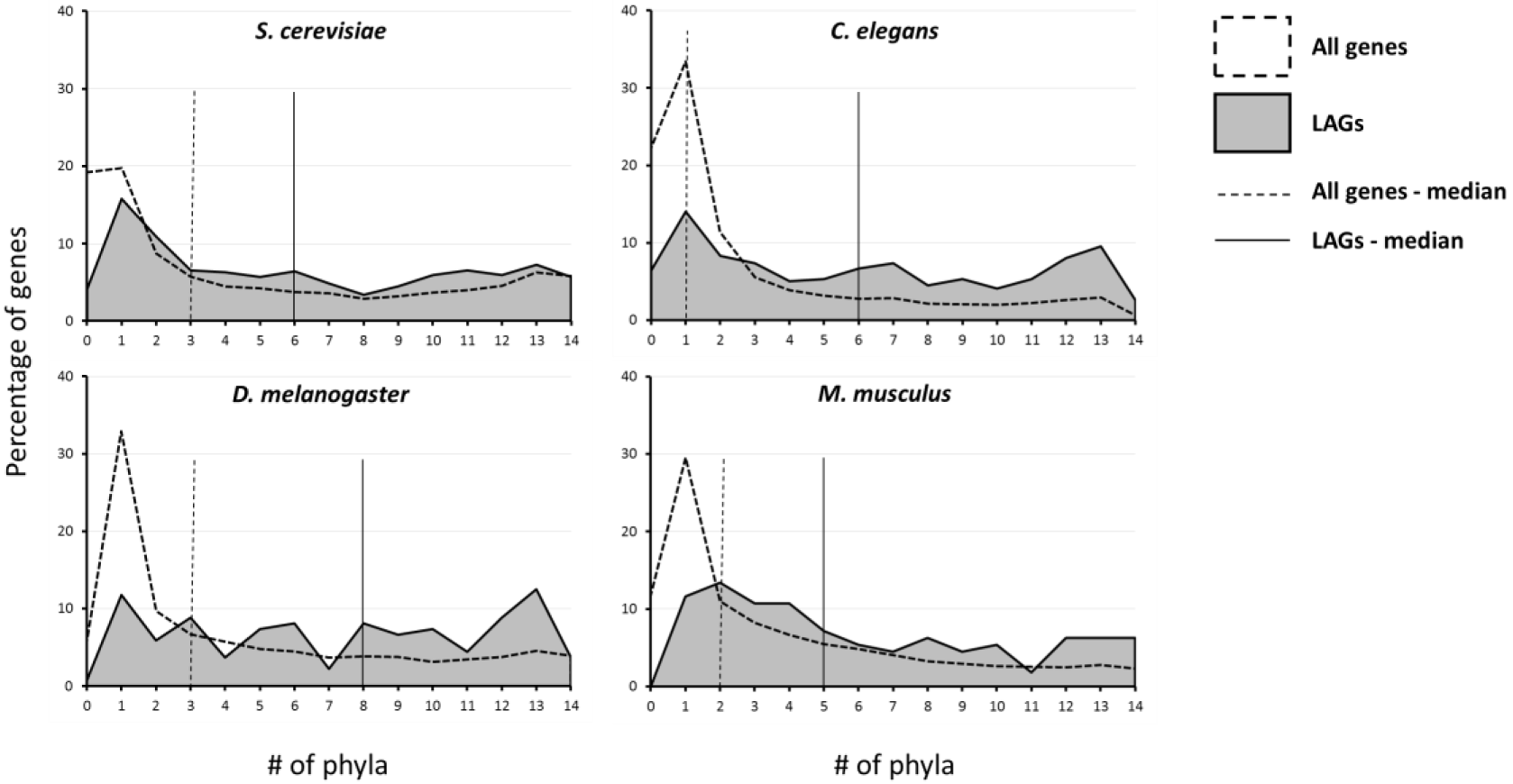
Distribution of LAGs according to the number of phyla in which LAGs have orthologs. Each graph represents the distribution of LAGs (grey area) discovered in the indicated model species. The entire proteome was used as a control (dotted line). X-axis depicts the number of phyla in which the genes have orthologs. The medians of the distributions are presented as vertical lines: dotted line for all genes and smooth black line for LAGs.

To get further insight into the universality of longevity-associated pathways, we carried out an enrichment analysis for LAGs that are conserved across a large number of phyla (“public”) and those that are specific to certain taxonomic group(s) (“private”). For the “private” analysis, we used the phylum of the corresponding model organim (as depicted in the Suppl. Table 5), since smaller taxonomic groups did not yield statistically conclusive results. Full detailed enrichment analysis is available in Suppl. Table 6.

Overall, the analysis of the most conserved (public) LAGs revealed that they fall under 4 major categories (Suppl. Tables 5 and 6): (i) Ribosome and translational processes; (ii) mitochondria and energy metabolism pathways (including the FoxO pathway); (iii) development, and (iv) DNA repair processes. These results provide a strong support for previous studies, in particular those by Smith et al. (2008), McElwee et al. (2007), and Freitas et al. (2011), and highlight the public role of these categories in the control of lifespan. It should however be noted that the number of LAGs was not always sufficient for a robust enrichment analysis, especially for the mouse and fly models (see Suppl. Table 6); the results from the yeast and worm models were more significant and thus more reliable.

Due to the high evolutionary conservation of LAGs, those that have orthologs only in the same phylum as the model species in which they were discovered are relatively small in number. Because of that, the enrichment analysis of these genes yielded less significant results (Suppl. Table 6). Nevertheless, the “private” list for *S. cerevisae* (i.e. yeast LAGs with orthologs only in Fungi) was found to be enriched with autophagy-related genes (Suppl. Table 5). For LAGs that have orthologs only in Nematoda, we surprisingly found enrichment in G-protein-related genes. Indeed, both autophagy and G-protein signaling represent basic and highly conserved processes which were shown to be involved in aging and longevity in various model organisms (Lans and Jansen 2007; Hahm et al. 2009; Rubinsztein et al. 2011; Schneider and Cuervo 2014). Yet, the unusual enrichment of these pathways in yeast and worms definitely highlights their importance in determination of longevity for these taxa specifically. For vertebrates, we found a significant enrichment in Neuroactive ligand-receptor interaction, which could reflect the importance of neuroendocrine regulation of aging and longevity in higher organisms (Dilman et al. 1986; Frolkis 1988; Blagosklonny 2013).

### Concordancy and discordancy in lifespan-modulating genetic interventions

Considering the conservation of many LAGs over a broad evolutionary distance, a valid question is whether modulating a given LAG in different species has a similar impact on longevity, i.e. lifespan extension or reduction. This was previously addressed for worm and yeast (Smith et al. 2008), where the genetic component of lifespan determination was found to be significantly conserved. Here, we broadened the question to *all* available model organisms. Namely, we compared all orthologs which were shown to have an impact on longevity in more than one species. Overall, we found that approximately 10% of LAGs’ orthologs (n = 184) were identified as such in at least two model organisms; 36 LAGs’ orthologs were identified in three and 20 in four model organisms. The number of concordant effects was significantly higher than the discordant ones (p < 0.003). That is, manipulation of LAGs has, more than often, the same effect in different species (Fig. 4, Suppl. Table 7). A substantial portion of the genetic interventions in yeast and worms could not be clearly defined as concordant or discordant with other model organisms (Fig. 4a, green), mostly due to a major difference in the methods of evaluation. In particular, in yeast studies, signs of premature aging and a reduction of lifespan could actually be just the results of reduced fitness but not mechanisms of aging *per se* (Tacutu et al. 2013).

**Fig. 4.**
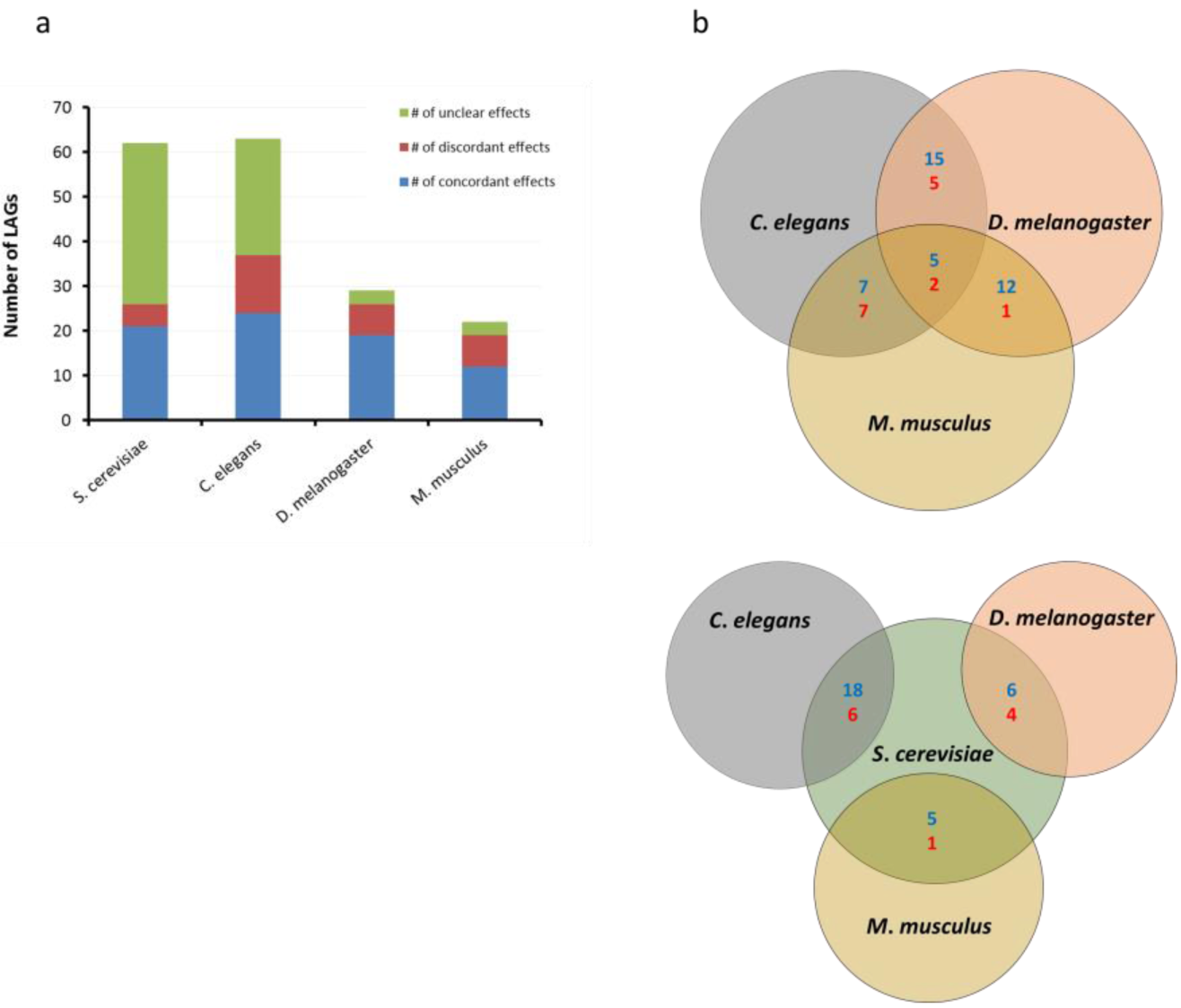
Concordancy in LAG manipulations across model organisms. Concordancy was determined according to the classification of LAGs as pro-or anti-longevity genes. That is, if a given LAG was determined as a pro-or anti-longevity gene in two or more species, it was termed “concordant”; otherwise, it was termed “discordant”. A detailed table is available in Suppl. Table 7. (**a**) Summary of the concordancy for LAGs from each model species which have also been tested in two or more species. (**b**) Venn diagram of the concordancy between species.

When looking at pairwise comparisons (Fig. 4B), it is evident that the level of concordancy is very high for some pairs of species (for example, *M. musculus* and *D. melanogaster*) and lower for others (for example, *M. musculus* and *C. elegans*). In order to discern what could bring about this difference, we calculated a conservation index for each pair of orthologs (as previously described (Huang et al. 2004)) and compared the results to the concordancy/discordancy of the effects. As seen in Suppl. Fig. 3, the observed discordancy could not be explained by sequence dissimilarity. One of the possible explanations for the observed discordancy is that in these cases orthologous LAGs were discovered by interventions which greatly differ from one another (e.g., knockout and overexpression). As such, if a given LAG is knocked-out and as a result the animal ages more rapidly, that LAG is defined as a “pro-longevity” gene; however, an overexpression of the same LAG will not necessarily increase lifespan. For example, a knockout of G protein, alpha subunit (*gpa-9*) in *C. elegans* increases maximum lifespan by up to 50%, but paradoxically, its overexpression also increases maximum lifespan (by 20%) (Schneider and Cuervo 2014). If such a difference occurs in the same species, more so could be expected when testing for effects on lifespan between different model organisms. Indeed, as is evident from Suppl. Fig. 4, the concordancy was significantly higher when a similar intervention was performed. Then, at least some of the discordancy could be explained by a variety in the methods of intervention.

Interestingly, only 5 LAGs (*Sod2*, *Sirt1*, *Mtor*, *Fxn*, and *Rps6kb1*; in total, 20 orthologs) were tested for their impact on longevity in all 4 model species. The manipulations of these genes showed a predominantly concordant effect on longevity, with the exception of *Fxn* (*Frh-1*) which has an opposite effect only in *C. elegans* (Suppl. Table 7). Altogether, the results indicate a clear trend of concordancy in the effects of LAG manipulations across model species despite a high evolutionary distance between them, and the observed cases of discordancy could mostly be attributed to technical rather than biological issues.

## Conclusions

Our wide-scale analysis of longevity-associated genes (LAGs) shows that their orthologs are consistently over-represented across diverse taxa, compared with the orthologs of other genes, and this conservation was relatively independent of evolutionary distance. The high evolutionary conservation was evident for LAGs discovered in all of the four major model organisms (yeast, *S. cerevisae*; worm, *C. elegans*; fly, *D. melanogaster*; mouse, *M. musculus*), but was especially relevant for *C. elegans*, where a large portion of LAGs were identified by genome-wide screens, thus minimizing potential biases. The results from the *C. elegans* analysis also suggest that antagonistic pleiotropy is a highly conserved principle of aging. Another important observation in our study was that the majority of manipulations on LAG orthologs in more than one model animal resulted in concordant effects on longevity. This strengthens the paradigm of “public” longevity pathways and of using model animals to study longevity, even across a large evolutionary distance. This notion is further strengthened when combined with the observation of Smith et al. (2008) who demonstrated that the existence of an ortholog is probably accompanied by a preserved role in longevity. Yet, we also observed LAGs that are highly conserved only in a limited number of taxa, or that displayed discordant effects when tested in more than one species, which could be attributed to “private” mechanisms of aging. Definitely, more comparative studies are warranted to better discriminate between private and public mechanisms, with unified methods of intervention and evaluation in mind. A recent study by Harel et al. (2015) could serve as a step in that direction by offering a new model of short-lived vertebrate species. In perspective, the combination of the existing data on LAGs with the emerging data on their expression throughout lifespan could bring about a deeper understanding of the role of genetic factors in aging and longevity.

## Methods

### Gene lists

#### Longevity-associated genes

The longevity-associated genes (LAGs) are defined as genes whose genetic manipulation in model organisms (*Mus musculus*, *Drosophila melanogaster*, *Caenorhabditis elegans* and *Saccharomyces cerevisiae*) was shown to significantly affect their lifespan. The list was obtained from Human Ageing Genomic Resources (HAGR) – GenAge database (http://genomics.senescence.info/genes/longevity.html; (Tacutu et al. 2013)).

#### Interactome genes

Interactome genes were extracted from the BioGrid database (http://www.thebiogrid.org; (Chatr-Aryamontri et al. 2015)) and were used as additional, high quality control for a more rigorous testing of the evolutionary conservation of LAGs.

#### Essential genes

Genes essential for the development and growth of *C. elegans* were extracted from WormBase (http://www.wormbase.org/ (Howe et al. 2016)).

### Determination of orthology

Ortholog determination for each gene was based on the InParanoid database - Eukaryotic Ortholog Groups (http://inparanoid.sbc.su.se/, (Sonnhammer et al. 2015)). The analysis was performed for 205 species (all species available excluding parasites; for a full list, see Suppl. Table 1). The ortholog extraction was performed automatically using software developed in our lab. The taxonomy of the species examined was based on the ITIS database (http://www.itis.gov/). The statistical significance of conservation for a group of genes was evaluated with the Chi-squared Goodness of fit test.

### Gene set enrichment

Enrichment analysis was performed using David Bioinformatics Resources 6.8 (https://david-d.ncifcrf.gov/ (Huang et al. 2009)), and WebGestalt (http://www.webgestalt.org/; (Wang et al. 2013)). The enrichment analysis was performed against 3 different backgrounds, including the whole genome, all LAGs, and the genes of the model organism under the same conservation criteria depicted in Suppl. Table 5.

## Declarations

### Authors’ contributions

All authors participated in data collection and analysis. In addition, TB wrote all the programs for data extraction from the databases used and, in particular, the ortholog extraction for large sets of genes and species and the calculation of the conservation index. HY wrote the manuscript and prepared the figures and tables. AB participated in writing the manuscript. RT was involved in the computational aspects of analysis. VEF coordinated the study.

## Acknowledgements

We would like to thank Prof. Marina Wolfson for her helpful suggestions and insightful comments.

## Supplementary Figures and Tables

**Suppl. Tables 1-4 and 6-7 are provided as additional Excel files.**

**Suppl. Table 5.**
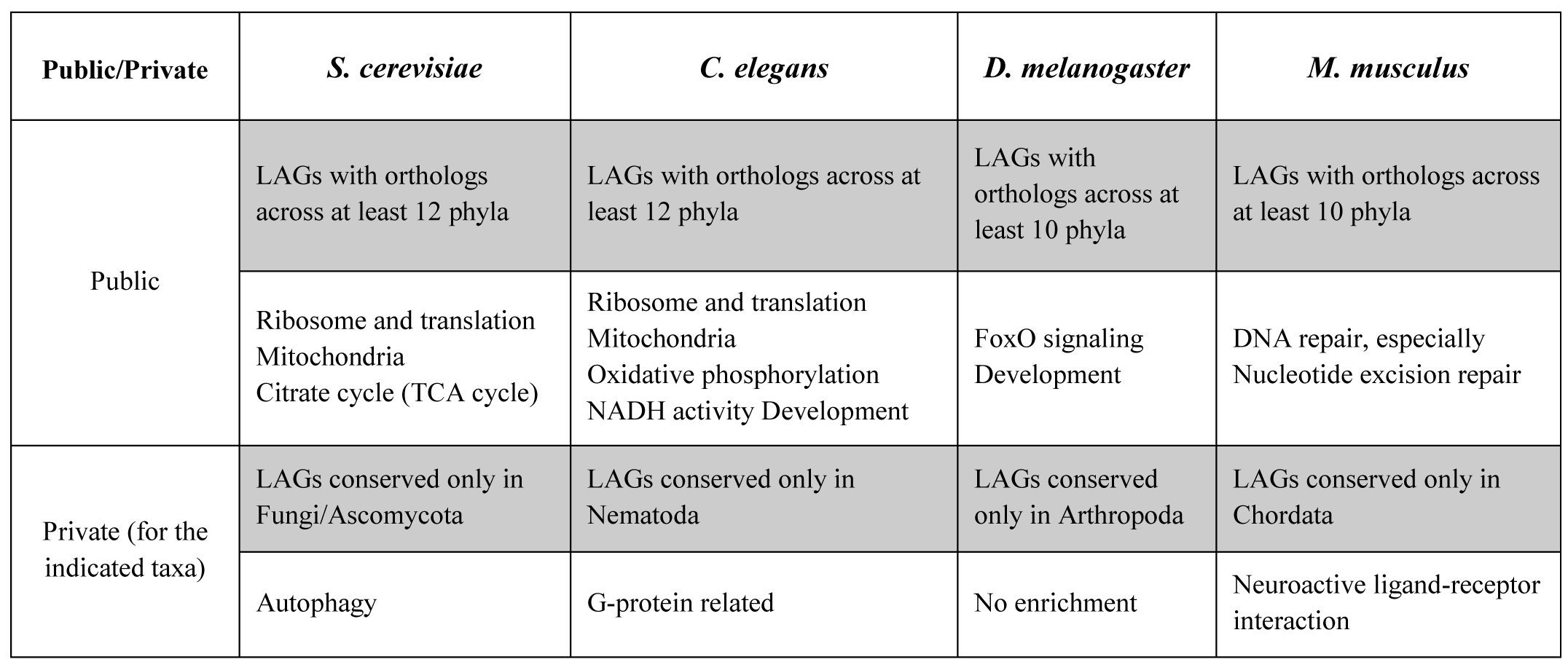
“Public” and “private” enriched categories. The table depicts the most enriched categories for lists of proteins of longevity-associated genes (LAGs) under different evolutionary conservation criteria (defined as the presence of orthologs across a listed number of phyla).

| Public/Private | *S. cerevisiae* | *C. elegans* | *D. melanogaster* | *M. musculus* |
| --- | --- | --- | --- | --- |
| Public | LAGs with orthologs across at least 12 phyla | LAGs with orthologs across at least 12 phyla | LAGs with orthologs across at least 10 phyla | LAGs with orthologs across at least 10 phyla |
|  | Ribosome and translation<br>Mitochondria<br>Citrate cycle (TCA cycle) | Ribosome and translation<br>Mitochondria<br>Oxidative phosphorylation<br>NADH activity Development | FoxO signaling<br>Development | DNA repair, especially<br>Nucleotide excision repair |
| Private (for the indicated taxa) | LAGs conserved only in Fungi/Ascomycota | LAGs conserved only in Nematoda | LAGs conserved only in Arthropoda | LAGs conserved only in Chordata |
|  | Autophagy | G-protein related | No enrichment | Neuroactive ligand-receptor interaction |

**Suppl. Fig. 1.**
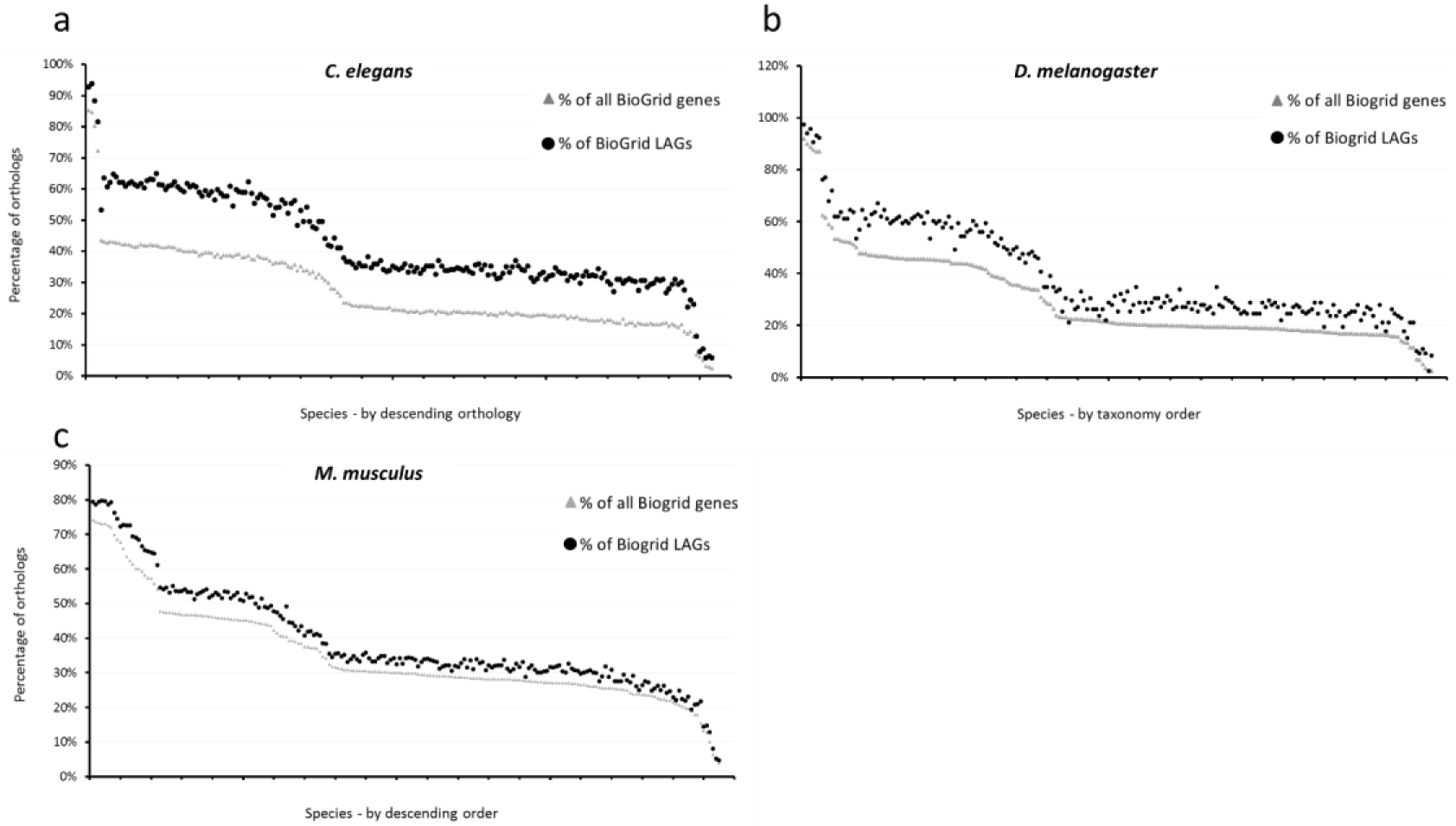
Percentage of Interactome LAG orthologs from the four model species. Each graph represents the LAGs discovered in the indicated model species. Each dot represents the percentage of orthologs between the model species and a different target species (total of 205 species from all kingdoms). Target species are ordered in descending order of orthology percentage as determined by the control. LAGs that are listed in BioGrid (black circle), entire proteome in BioGrid (grey triangle). **(a)** *C. elegans*, n = 3,865 for control and 343 for LAGs; **(b)** D. *melanogaster*, n = 8,026 for control and 118 for LAGs; **(c)** S. *cerevisiae*, n = 5,783 for control and 823 for LAGs.

**Suppl. Fig. 2.**
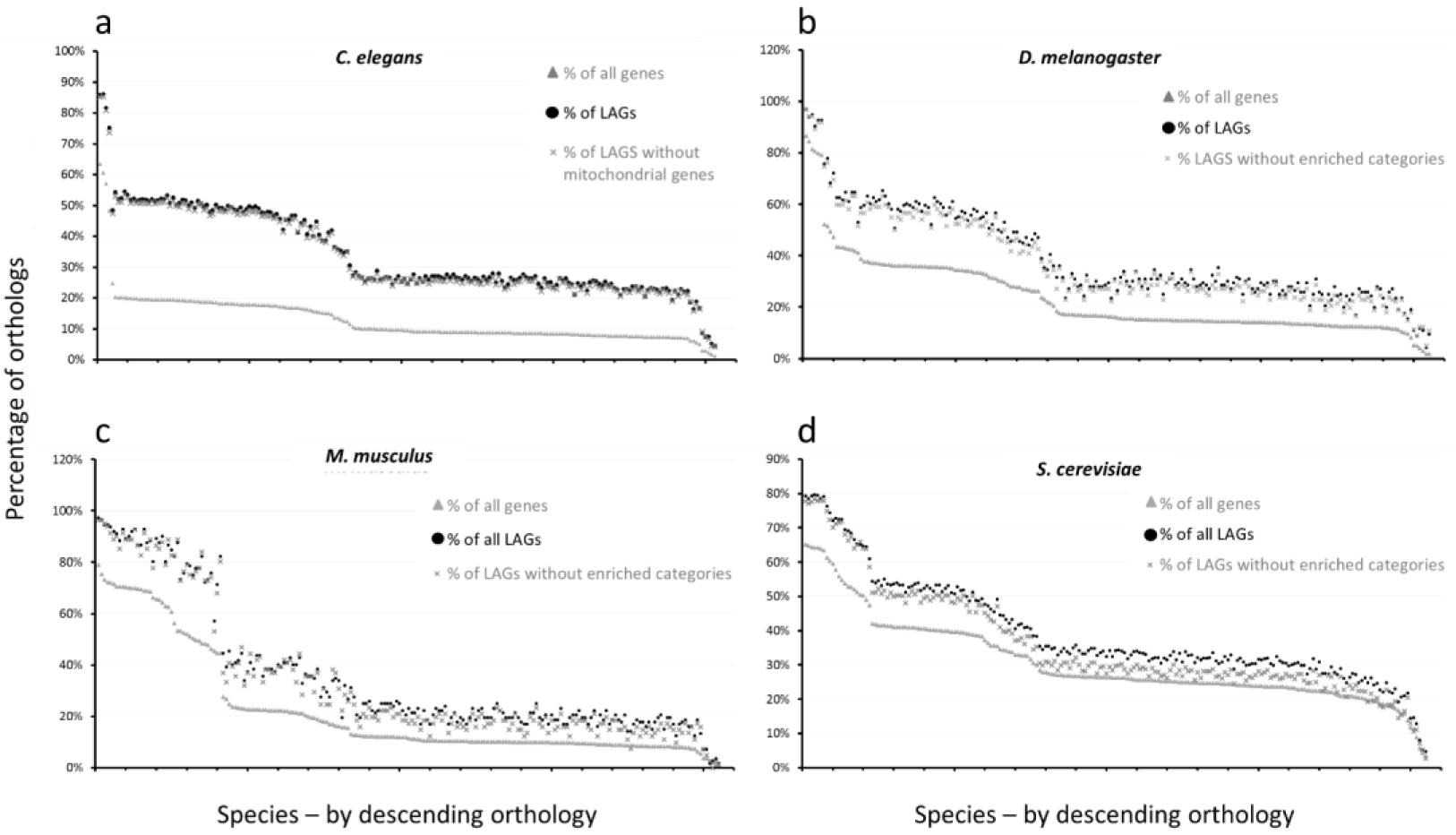
Percentage of LAG orthologs from the four model species after exclusion of proteins from enriched categories. Each graph represents the LAGs discovered in the indicated model species. Each dot represents the percentage of orthologs between the model species and a different target species (total of 205 species from all kingdoms). Target species are ordered in descending order of orthology percentage as determined by the control. LAGs (black circle), entire proteome (grey triangle) and LAGs after exclusion of proteins in enriched categories (grey x). (**a)** *C. elegans*, n = 20,325 for control, 733 for LAGs and 689 LAGs without mitochondrial genes; (**b)** *D. melanogaster*, n = 13,250 for control, 136 for LAGs and 122 for LAGs excluding enriched categories; (**c)** *M. musculus*, n = 21,895 for control,112 for LAGs and 73 LAGs excluding enriched categories; (**d)** *S. cerevisiae*, n = 6,590 for control, 824 for LAGs and 551 LAGs excluding enriched categories. All differences presented are highly significant (p < 10^-10^).

**Suppl. Fig. 3.**
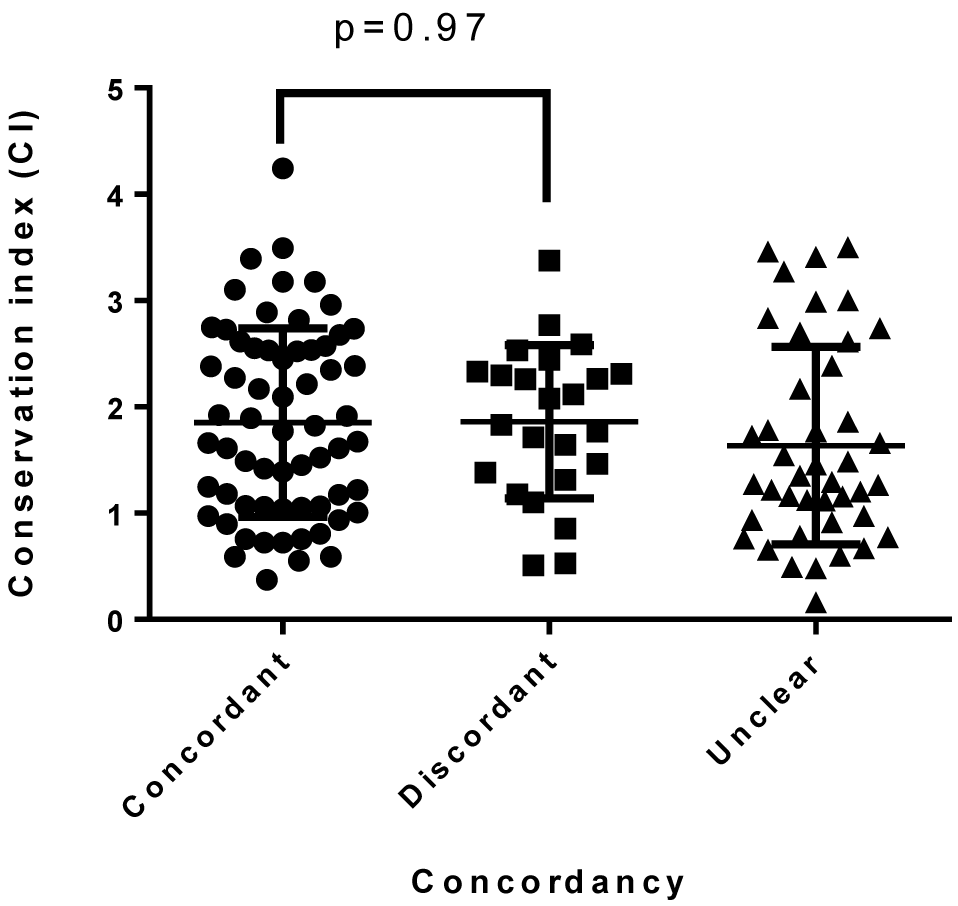
Conservation index (CI) compared to concordancy of longevity effects. Each dot represents a pairwise comparison between orthologs of LAGs that were tested for their effect on longevity in more than one model species. The conservation index is the pairwise alignment score normalized to the protein amino acid length.

**Suppl. Fig. 4.**
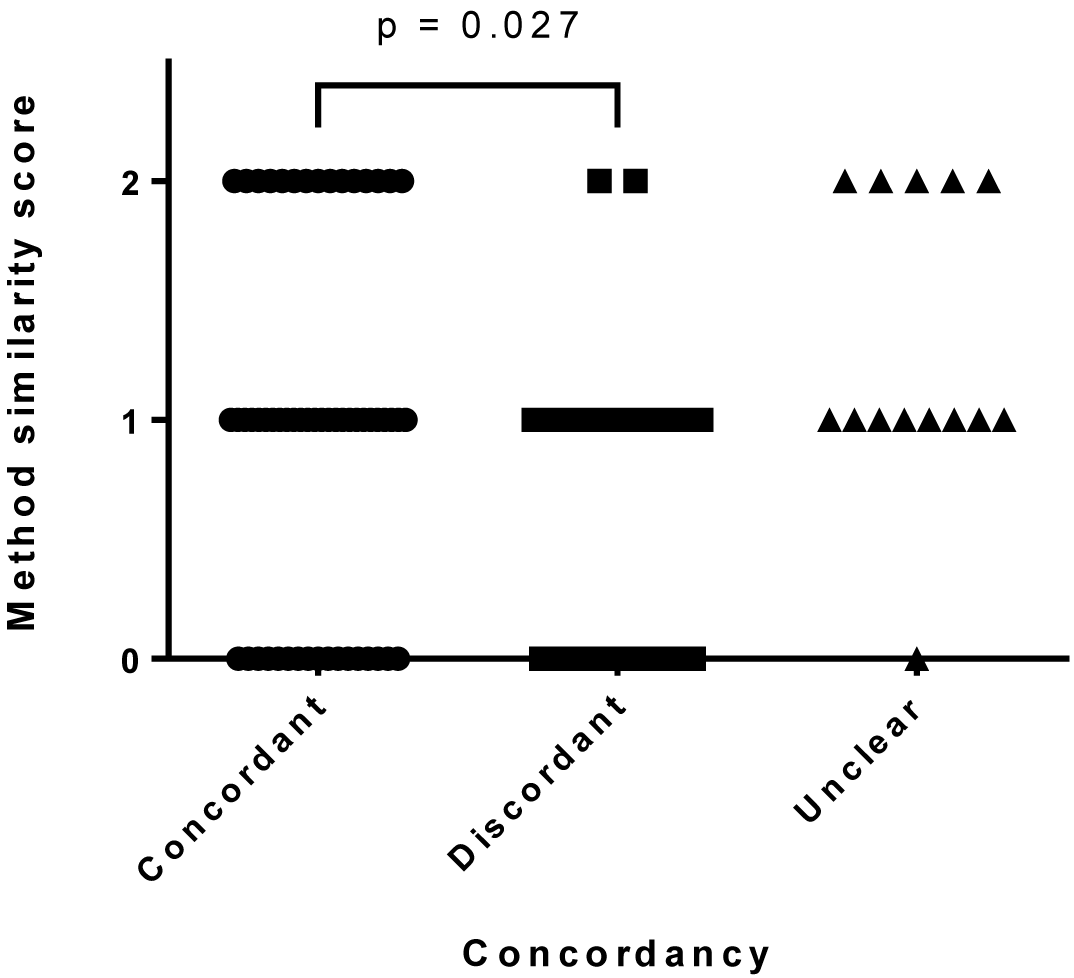
Method similarity score compared to concordancy of longevity effects. Each dot represents a pairwise comparison between orthologs of LAGs that were tested for their effect on longevity in more than one model species. The method similarity score was determined as: 0 = interventions of opposite directions (e.g. knockout and overexpression); 1 = intervention of the same direction but with varied methods (e.g. knockout and RNAi); 2 = interventions that are identical or very close to identical (e.g. knockout and knockout).

